# The Lands cycle modulates plasma membrane lipid organization and insulin sensitivity in skeletal muscle

**DOI:** 10.1101/2019.12.23.887232

**Authors:** Patrick J. Ferrara, Xin Rong, J. Alan Maschek, Anthony R.P. Verkerke, Piyarat Siripoksup, Haowei Song, Karthickeyan C. Krishnan, Jordan M. Johnson, John Turk, Joseph A. Houmard, Aldons J. Lusis, James E. Cox, Saame Raza Shaikh, Peter Tontonoz, Katsuhiko Funai

## Abstract

Aberrant lipid metabolism promotes the development of skeletal muscle insulin resistance, but the exact identity of lipid-mediated mechanisms relevant to human obesity remains unclear. A comprehensive lipidomic analyses of primary myocytes from lean insulin-sensitive (LN) and obese insulin-resistant (OB) individuals revealed several species of lysophospholipids (lyso-PL) that were differentially-abundant. These changes coincided with greater expression of lysophosphatidylcholine acyltransferase 3 (LPCAT3), an enzyme involved in phospholipid transacylation (Lands cycle). Strikingly, mice with skeletal muscle-specific knockout of LPCAT3 (LPCAT3-MKO) exhibited greater muscle lyso-PC/PC, concomitant with greater insulin sensitivity *in vivo* and insulin-stimulated skeletal muscle glucose uptake *ex vivo*. Absence of LPCAT3 reduced phospholipid packing of the cellular membranes and increased plasma membrane lipid clustering, suggesting that LPCAT3 affects insulin receptor phosphorylation by modulating plasma membrane lipid organization. In conclusion, obesity accelerates the skeletal muscle Lands cycle, whose consequence might induce the disruption of plasma membrane organization that suppresses muscle insulin action.

## Introduction

Type 2 diabetes is the 7^th^ leading cause of death in the United States (1) and is a major risk factor for cardiovascular disease, the leading cause of death (2). Skeletal muscle is the site of the largest glucose disposal in humans (3, 4). Insulin resistance in skeletal muscle is a necessary precursor to type 2 diabetes (5) and can be triggered by aberrant lipid metabolism (6–8). Several classes of lipids have been implicated in initiating cellular signals that suppress insulin action, but there has not been a clear consensus that these molecules are upregulated in skeletal muscle insulin resistance that occurs in the human population (9–14).

A difficulty in accurately measuring the muscle lipidome is confounded by the intramyofibrillar adipocytes which are particularly abundant in muscle biopsy samples from obese humans. Human skeletal muscle cells (HSkMC) are primary myoblasts that can be isolated, propagated, and differentiated from muscle biopsies. This *in vitro* system provides a unique model to study the skeletal muscle lipidome and signaling pathways free of contaminating cell types and circulating factors that affect muscle metabolism. Importantly, these HSkMC are known to retain their insulin sensitivity phenotype *ex vivo*, providing a platform to study mechanisms directly relevant to human physiology (15, 16).

In this study, we harvested HSkMC from lean insulin-sensitive (LN) and obese insulin-resistant (OB) subjects (Subject Characteristics: Table S1). We then propagated and differentiated these samples for analyses of the muscle lipidome, gene expression profile, insulin signaling, and membrane properties (Figure 1A). This approach led us to examine the lysophospholipid (lyso-PL) remodeling pathway (Lands cycle) as a potential diet-responsive mechanism that regulates skeletal muscle insulin action. Below we provide evidence that implicates this pathway in the pathogenesis of diet-induced skeletal muscle insulin resistance. Genetic or pharmacologic suppression of this pathway was sufficient to enhance skeletal muscle insulin action *in vitro* and *in vivo*.

**Figure 1:**
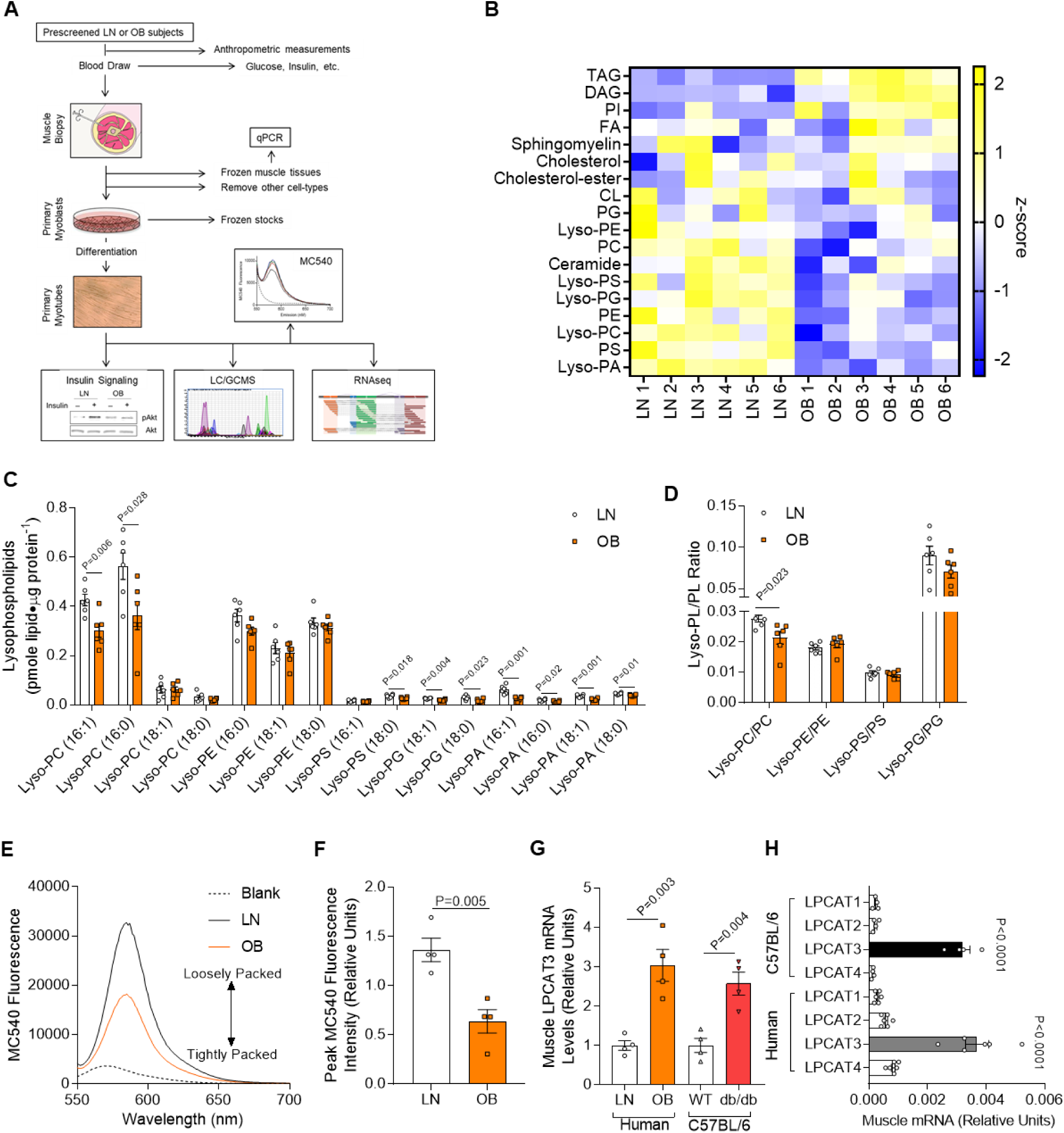
Lipidomic analyses of skeletal muscle samples from lean and obese human subjects. (A) A schematic of the workflow for the clinical study. (B-D) Lipidomic analysis of HSkMC from lean insulin-sensitive (LN) and obese insulin-resistant (OB) subjects. (B) Heat map of lipid content by class. (C) Species of lysophospholipids. (D) Lysophospholipid to phospholipid ratio (*n*=6). (E&F) Quantification of MC540 fluorescence in LN and OB HSkMC (*n*=4). (G) LPCAT3 mRNA in muscle biopsies from LN or OB subjects (left), and in skeletal muscle of wild type (WT) or a db/db (right) mice (*n*=4). (H) Expression of all isoforms of LPCAT in muscle samples from mouse (*n*=4) or human (*n*=6) skeletal muscle. (C,D,F&G) Two-tailed t-tests. (H) One-way ANOVA followed by post-hoc multiple comparisons. All data are represented as mean ± SEM.

## Results

A global lipidomic analysis of LN and OB myotubes revealed many classes of lipids that were differentially abundant (Figure 1B, Figure S1A-L). Among these, several species of lysophospholipids (lyso-PL), intermediates of the Lands cycle (17), were lower in OB HSkMC compared to LN (Figure 1C), a finding not previously described for an insulin-resistant state. While species of many classes of lyso-PL were reduced with obesity, the ratio of lyso-PL to its parent phospholipid were significantly lower for only lysophosphatidylcholine (lyso-PC)/phosphatidylcholine (PC) (Figure 1D). Previous studies suggest that altering the lyso-PC content of cell membranes is sufficient to alter the physical properties of membranes (18, 19). Consistent with this notion, phospholipid packing of LN and OB myotubes were remarkably different, with OB cells exhibiting more tightly packed membrane head groups compared to LN (Figure 1E&F). This occurred in the absence of changes in the phospholipid acyl-chain saturation index (Figure S1M).

What is the molecular mechanism by which obesity promotes a lower abundance of lyso-PL in skeletal muscle? LN and OB HSkMC utilized for the lipidomic analyses were cultured *ex vivo* for several weeks in identical media conditions. Thus, differences in the lipidome of these samples are likely the result of genetic and/or epigenetic influences, instead of hormonal or neuronal inputs that alter cells *in vivo*. We reasoned that such differential programming might be expected to manifest in gene expression profiles. A whole transcriptome sequencing of LN and OB myotubes revealed that lyso-PC acyltransferase 3 (LPCAT3), an enzyme of the Lands cycle, was more highly expressed with obesity. These findings were recapitulated in muscle biopsy samples (*not* myotubes) from LN and OB individuals as well as muscle tissue from wild-type and db/db mice (Figure 1G). The Lands cycle represents a series of phospholipid-remodeling reactions by which acyl-chains become transacylated (17). Of the thirteen lyso-PL acyltransferase enzymes (20), LPCAT3 has the highest affinity for 16:0 and 18:0 lyso-PC, consistent with the specificity of reduced lyso-PC/PC (21, 22). Silencing of LPCAT3 in fibroblasts has been shown to increase Akt phosphorylation (23), while incubation of the same cells with 16:0/20:4 PC decreased Akt phosphorylation due to plasma-membrane specific effects (24). Mice with a liver-specific deletion of LPCAT3 exhibit enhanced ordering of membranes (25). In both human and mouse skeletal muscle, LPCAT3 is very highly expressed compared to other isoforms of LPCAT (Figure 1H), and skeletal muscle LPCAT3 expression is directly correlated to circulating insulin in 106 mouse strains (data not shown, bicor=0.296, P=0.0016) (26).

To study the role of LPCAT3 on skeletal muscle insulin action, we performed a lentivirus-mediated shRNA knockdown of LPCAT3 (Scrambled, SC; LPCAT3 knockdown, KD) in C2C12 myotubes (Figure 2A). LPCAT3 knockdown did not affect protein content for MyoD, various MHC isoforms, and respiratory complexes (Figure S2A-C), suggesting that the deletion of LPCAT3 has no effect on myotube lineage or mitochondrial density. Targeted lipidomic analyses revealed that LPCAT3 knockdown increased lyso-PC and decreased PC (Figure 2B&C), substrates and products of the LPCAT3-mediated reaction, respectively (27, 28). Together, these differences were sufficient to elevate lyso-PC/PC with LPCAT3 deletion (Figure 2D). Similar effects were seen with lipid species composed of an ethanolamine head group (Figure S2D-F), while the phospholipid saturation index increased with LPCAT3 knockdown (Figure S2G). Analogous to differences observed in LN and OB HSkMC, LPCAT3 deletion reduced phospholipid head group packing (Figure 2E&F). We then incubated SC and KD cells in a submaximal concentration of insulin to assess insulin signaling events. Strikingly, inhibition of LPCAT3 robustly enhanced insulin signaling with or without insulin (Figure 2G, Figure S2H). Notably, the increase occurred at the level of the insulin receptor (IR), a node that is localized in the phospholipid-rich plasma membrane. Consequently, LPCAT3 deletion enhanced insulin-stimulated glycogen synthesis (Figure 2H), suggesting that this intervention increases skeletal muscle insulin sensitivity *in vitro* (due to low GLUT4:GLUT1 stoichiometry, insulin-stimulated glucose uptake is not an ideal surrogate for insulin sensitivity in C2C12 myotubes). LPCAT3 knockdown also enhanced insulin signaling in HSkMC from obese subjects (Figure 2G).

**Figure 2:**
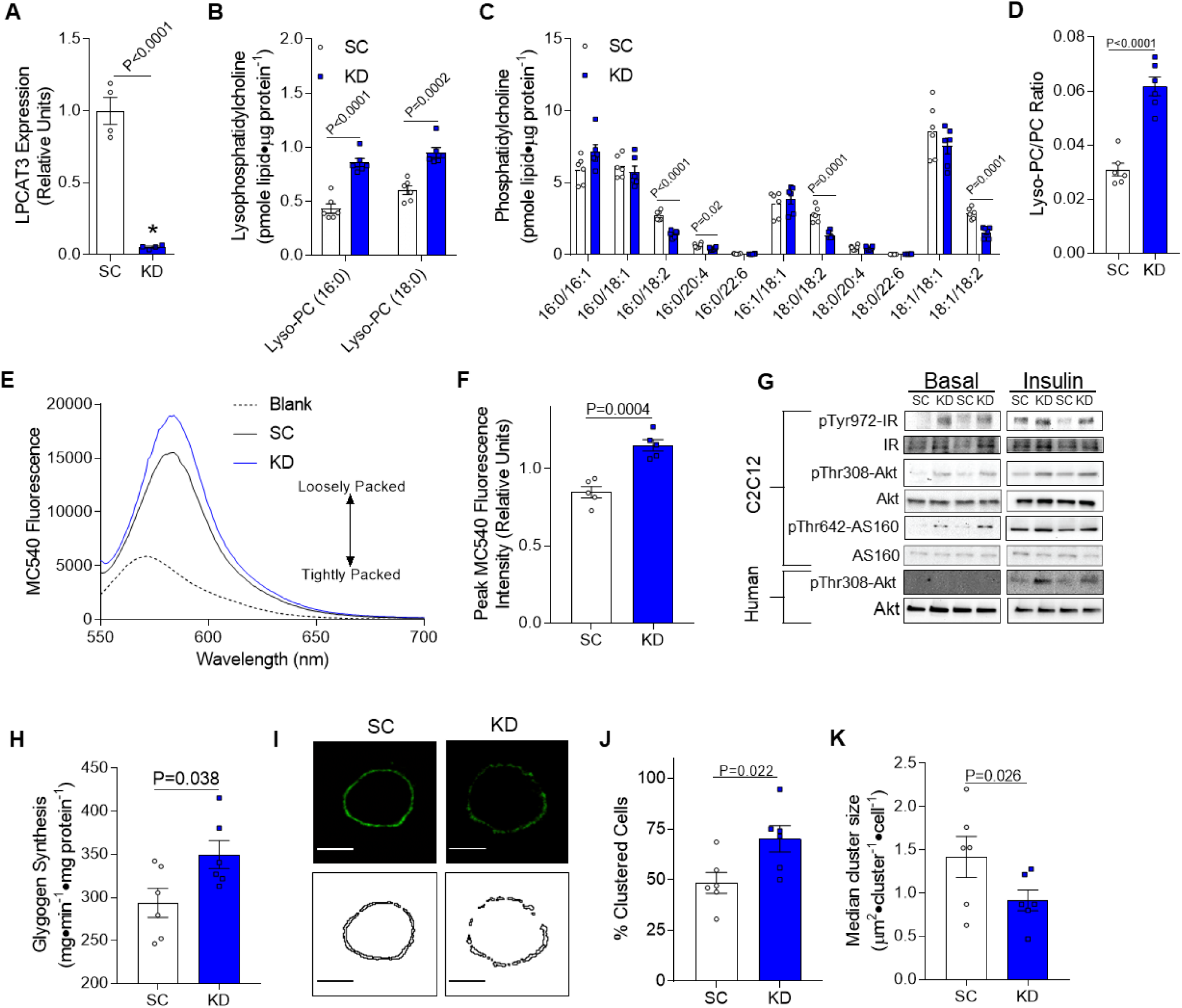
LPCAT3 knockdown enhances skeletal muscle insulin sensitivity *in vitro*. (A) LPCAT3 mRNA levels in myoblasts infected with lentiviruses expressing shRNA for scrambled (SC) or LPCAT3 sequences (KD) and differentiated into myotubes (*n*=4). (B-D) Lipids were extracted from C2C12 myotubes for analysis between SC and KD cells. Quantification of (B) lysophosphatidylcholine (lyso-PC), (C) phosphatidylcholine (PC), and (D) total lyso-PC/PC (*n*=6). (E&F) Quantification of MC540 fluorescence in SC and KD myotubes (*n*=5). (G) Phosphorylation and total protein of IR, Akt, and AS160 were measured via Western blot with (0.6 nM) or without insulin in C2C12 myotubes (top) and human primary skeletal muscle cells (bottom). (H) Glycogen synthesis was quantified in C2C12 cells incubated with insulin (12 nM) (*n*=6). (I-K) GM-1 enriched microdomains were labeled in SC and KD round-up myotubes. (I) Plasma membrane GM-1 localization was visualized (top panels: fluorescence images, bottom panels: binary images). (J) Cells were scored as clustered or non-clustered between SC and KD myotubes. (K) Particle size was measured for each cell in 6 separate experiments and the median for each experiment was used as a representative of that experiment (n=35-50/experiment, 6 separate experiments). (A-D,F,H,J&K) Two-tailed t-tests were performed. All data are represented as mean ± SEM.

The organization and clustering of plasma membrane microdomains is linked to the induction of tyrosine-kinase signaling events, such as IR signaling (29–31). Because LPCAT3 deletion promoted enhanced insulin signaling at the level of IR phosphorylation, we visualized the organization of plasma membrane microdomains with labeling and patching of plasma membrane GM-1, a known marker of microdomains. Indeed, a greater proportion of C2C12 cells with LPCAT3 knockdown exhibited clustering of GM-1 enriched microdomains (Figure 2I, top &J). Furthermore, LPCAT3 deletion decreased the size of these clusters (Figure 2I, bottom &K), with no differences in total fluorescence from each cell (Figure S2I). These data indicate that LPCAT3 inhibition induces a reorganization of plasma membrane microdomains, potentially explaining increased IR phosphorylation.

Next, we examined whether inhibition of muscle LPCAT3 would promote greater insulin sensitivity *in vivo*. Mice with tamoxifen-inducible skeletal muscle-specific knock-out of LPCAT3 (LPCAT3-MKO) were generated by crossing the HSA-MerCreMer mice (34) with LPCAT3 conditional knock-out mice (exon3 of the *Lpcat3* gene flanked with *loxP* sites) (25) (Figure 3A). This strategy successfully yielded mice with suppressed LPCAT3 expression in skeletal muscle without affecting other tissues (Figure 3B), and without compensatory upregulation of other members of the LPCAT family (Figure 3C). Control and LPCAT3-MKO mice gained weight equally when fed a high-fat diet (HFD, Figure 3D) with no difference in adipose tissue weight at the end of diet intervention (Figure 3E). Food consumption, whole-body oxygen consumption, spontaneous activity, and respiratory exchange ratio were similarly not different between groups (Figure 3F-I). Fasting glucose, insulin, and glucose tolerance (Figure 3J-L) were unchanged, but circulating insulin during the glucose tolerance test was substantially lower in LPCAT3-MKO mice compared to the control group (Figure 3M).

**Figure 3:**
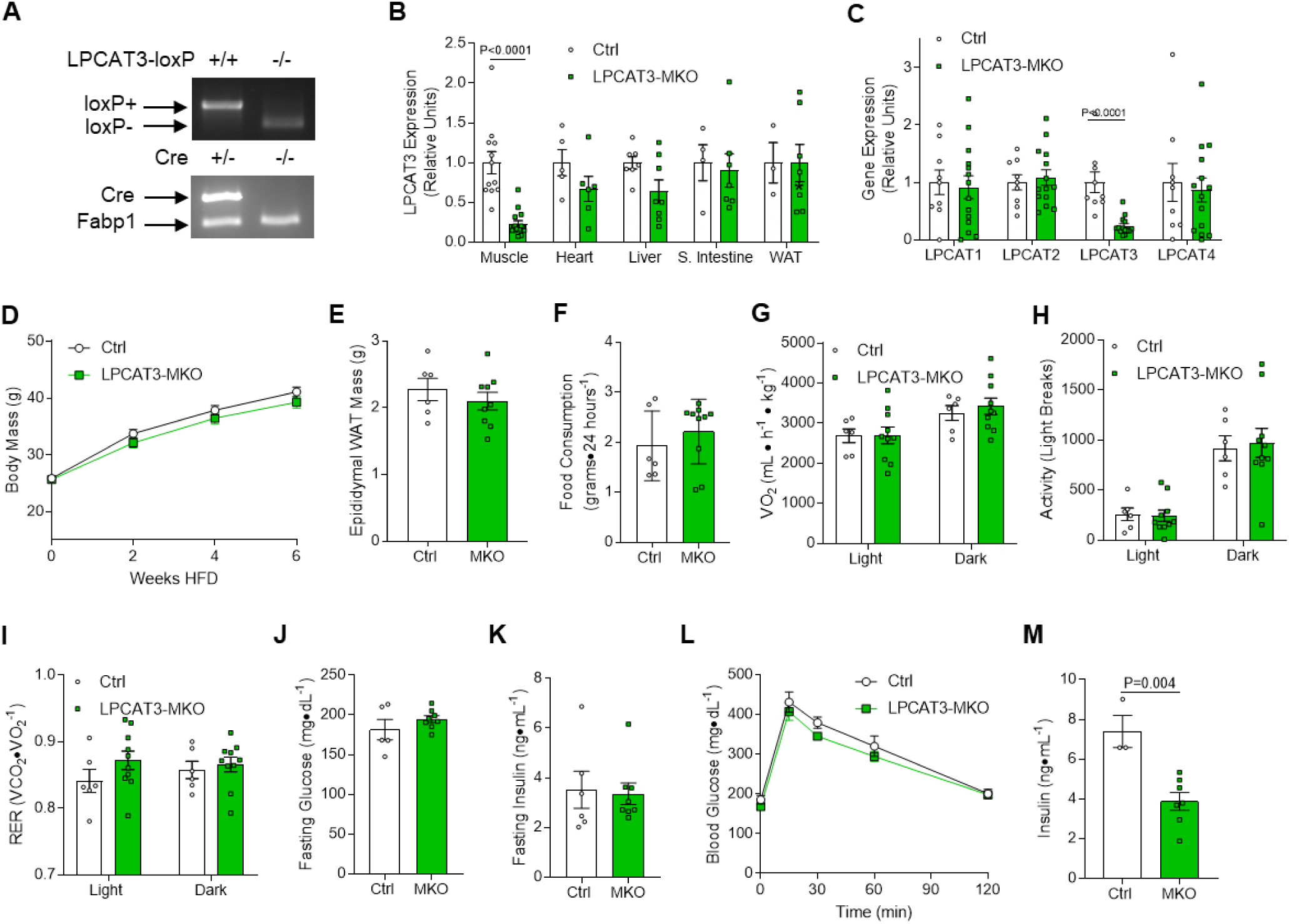
Whole-body phenotyping of LPCAT3-MKO mice. (A) Mice with tamoxifen-inducible skeletal muscle-specific Cre-recombinase (HSA-MerCreMer+/-) were crossed with mice with *LoxP* sites flanking exon3 of the *Lpcat3* gene (LPCAT3cKO+/+) to generate skeletal muscle-specific inducible knock out of LPCAT3 (LPCAT3cKO+/+, HSA-MerCreMer+/-) (LPCAT3-MKO). Littermates (LPCAT3cKO+/+, HSA-MerCreMer-/-) (Ctrl) were used as control mice for all experiments. (B) LPCAT3 mRNA in tibialis anterior (TA, Muscle), heart, liver, small intestine (S. Intestine), and inguinal white adipose tissue (WAT) (muscle: Ctrl *n*=12, MKO *n*=15; heart: Ctrl *n*=5, MKO *n*=6; liver: Ctrl *n*=7, MKO *n*=8; S. Intestine: Ctrl *n*=4, MKO *n*=7; WAT: Ctrl *n*=3, MKO *n*=7). (C) mRNA of all LPCAT isoforms in TA muscles of Ctrl and LPCAT3-MKO mice (Ctrl *n*=9, MKO *n*=14) (D) Body mass during high-fat diet (HFD) feeding in Ctrl and LPCAT3-MKO mice (Ctrl *n*=8, MKO *n*=11). (E) Epididymal WAT mass (Ctrl *n*=6, MKO *n*=9). (F-I) Ctrl and LPCAT3-MKO mice were placed in metabolic chambers for measurement of (F) food consumption, (G) VO_2_, (H) activity, and (I) respiratory exchange ratio (RER) (Ctrl *n*=6, MKO *n*=10). (J) Fasting glucose (Ctrl *n*=5, MKO *n*=9). (K) Fasting insulin (Ctrl *n*=6, MKO *n*=9). (L) Intraperitoneal glucose tolerance test (Ctrl *n*=6, MKO *n*=8). (M) Serum insulin at the 30-minute time point of the glucose tolerance test (Ctrl *n*=3, MKO *n*=8). All data except (A) are from HFD-fed mice. (B,C,E-K&M) Two-tailed t-tests or (D&L) 2-way ANOVA with Sidak’s multiple comparisons test were performed. All data are represented as mean ± SEM.

To evaluate whether improved glycemic efficiency was attributable to greater skeletal muscle insulin sensitivity, we quantified insulin-stimulated skeletal muscle glucose uptake *ex vivo*. Isolated muscles from HFD-fed control and LPCAT3-MKO mice were incubated with or without a submaximal concentration of insulin for the measurement of 2-deoxy-glucose uptake. Indeed, insulin-stimulated skeletal muscle glucose uptake was robustly enhanced in LPCAT3-MKO mice compared to control (Figure 4A). The increase in glucose uptake coincided with augmented insulin-stimulated Akt phosphorylation (Figure 4B&C), similar to C2C12 and human primary myotubes (Figure 2). These results suggest that inhibition of muscle LPCAT3 increases skeletal muscle insulin sensitivity *in vivo*. Similar to results from LPCAT3 knockdown *in vitro*, muscles from LPCAT3-MKO mice had elevated lyso-PC (16:0 and 18:0) (Figure 4D) and lower levels of PC species known to be the main products of the LPCAT3 reaction (16:0/18:2 and 16:0/22:4) (Figure 4E) (22, 35, 36). As a result, lyso-PC/PC was ∼2-fold greater in LPCAT3-MKO mice compared to control (Figure 4F). In contrast, lyso-PE/PE or phospholipid saturation index was unaltered between control and LPCAT3-MKO muscles (Figure S3A-D), similar to the lipidome in LN and OB HSkMC (Figure 1&S1). Muscles from control and LPCAT3-MKO did not differ in mass, length, force-generating capacity, fiber-type distribution, or content of proteins in the electron transport chain (Figure 4G-L, Figure S3E-G).

**Figure 4:**
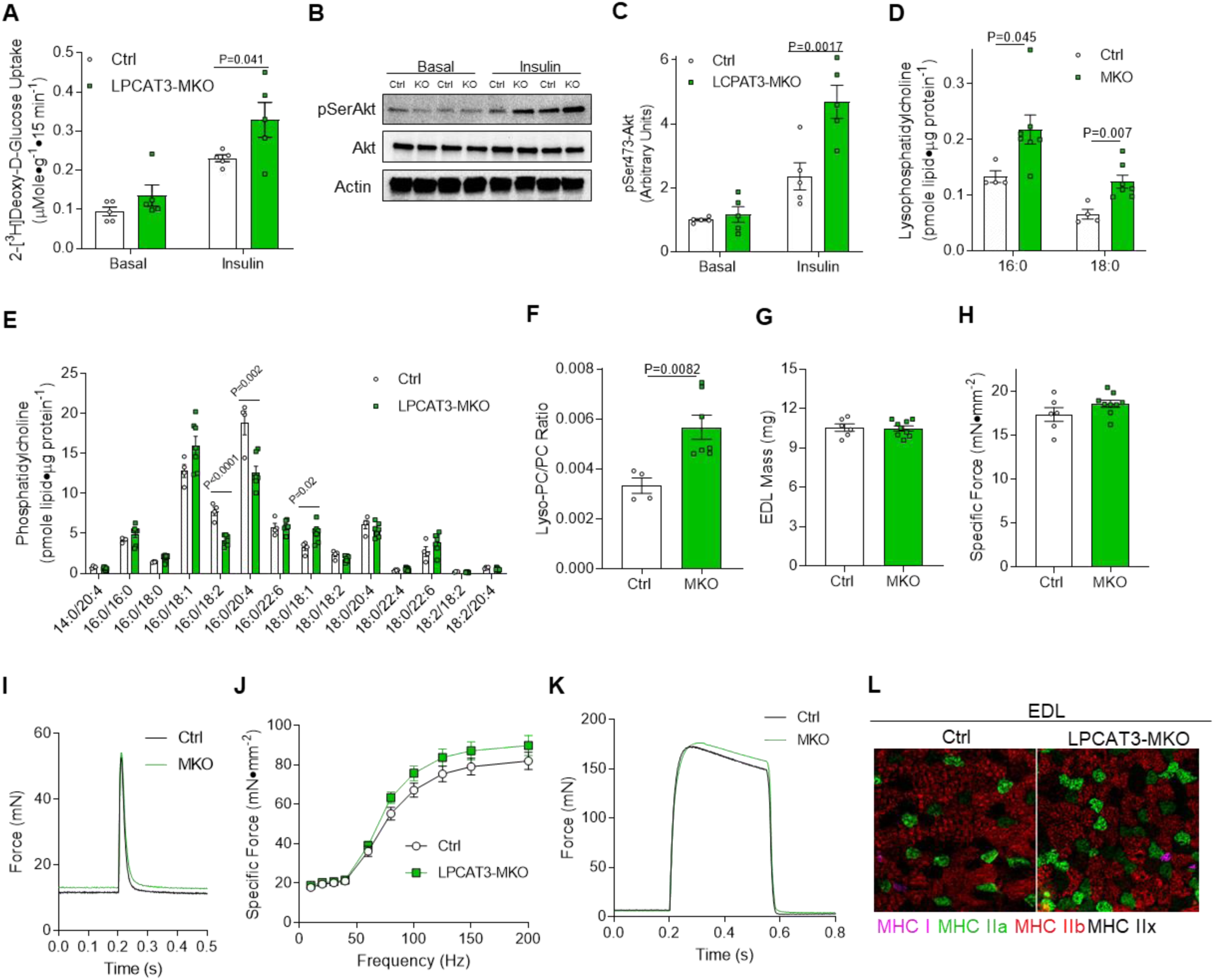
LPCAT3-MKO mice are protected from diet-induced skeletal muscle insulin resistance. (A-C) Soleus muscles were dissected and incubated with or without 200 µU/mL of insulin. (A) *Ex vivo* 2-deoxyglucose uptake (*n*=5). (B&C) Ser473 phosphorylation and total Akt (Ctrl *n*=5, MKO *n*=6). (D-F) Lipids were extracted from gastrocnemius muscles of Ctrl and LPCAT3-MKO mice for mass spectrometric analysis. Quantification of (D) lyso-PC, (E) PC, and (F) total lyso-PC/PC (Ctrl *n*=4, MKO *n*=7). (G-L) Extensor digitorum longus (EDL) muscles of Ctrl and LPCAT3-MKO mice were dissected for measurement of (G) mass, (H&I) force produced with a pulse stimulation, (J&K) force produced with tetanic stimulation ranging from 10-200 Hz (K, force tracing at 200 Hz stimulation) (Ctrl *n*=6, MKO *n*=9), and (L) skeletal muscle fiber-type (MHC I: pink, MHC IIa: green, MHC IIb:red, and MHC IIx: negative). All data are from HFD-fed mice. (A,C&J) 2-way ANOVA with Sidak’s multiple comparisons test or (D-H) two-tailed t-tests were performed. All data are represented as mean ± SEM.

How does the inhibition of the Lands cycle promote greater insulin action in skeletal muscle? LPCAT3 deficiency enhanced insulin signaling at the level of IR which was concomitant with altered plasma membrane lipid organization (Figure 2), suggesting that changes in plasma membrane properties may mediate the insulin-sensitizing effects. Membrane organization is vital to insulin action, as IR is localized to highly ordered microdomains on the plasma membrane (37, 38). The interaction between caveolae and IR enhances insulin signaling in other cell types (39, 40). Mice that lack caveolin-3 (cav3), a skeletal muscle-specific scaffolding protein critical in the formation of caveolae on the plasma membrane, exhibit skeletal muscle insulin resistance due to plasma membrane-specific effects on the IR (41, 42). Overexpression of dominant-negative cav3 leads to decreased glucose uptake and glycogen synthesis in C2C12 cells, which is attributed to decreased Akt phosphorylation (43–45). Conversely, an increase in wildtype cav3 expression is sufficient to enhance Akt phosphorylation and glucose uptake (46). Indeed, LPCAT3 knockdown substantially increased cav3 content in C2C12 myotubes (Figure 5A). To examine the possibility that the absence of LPCAT3 increases the abundance of lipids in caveolae, we isolated membrane fractions from C2C12 myotubes with or without LPCAT3 deletion and subjected them for further purification by density-gradient ultracentrifugation.

**Figure 5:**
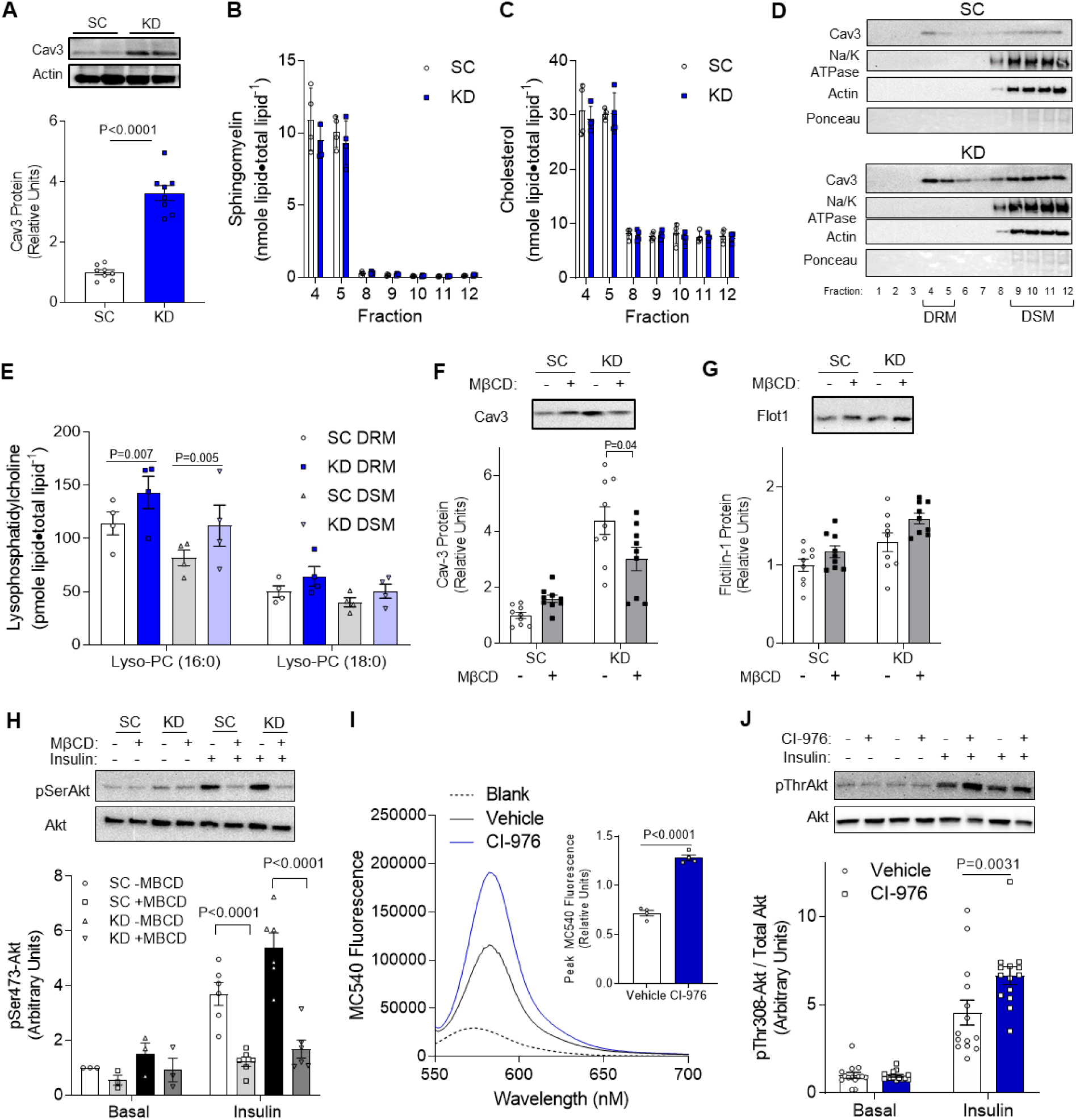
LPCAT3 deletion alters plasma membrane organization. C2C12 cells were infected with lentiviruses expressing shRNA for scrambled (SC) or LPCAT3 (KD) and differentiated into myotubes. (A) Caveolin-3 (cav3) protein content. (B&C) Detergent-resistant membranes (DRM; fractions 4&5) and detergent-soluble membranes (DSM; fractions 8-12) were isolated and lipids were extracted for quantification of (B) sphingomyelin and (C) cholesterol (*n*=4). (D) Cav3, Na/K ATPase, actin, and total protein content were assessed via Western blot in all fractions from the sucrose gradient. (E) Lyso-PC levels in DSM and DRM isolations (*n*=4). (F-H) C2C12 myotubes were incubated in the presence (10 mM) or absence of methyl-beta-cyclodextrin (MβCD) for 1 hour. (F&G) MβCD successfully depletes cav3 (P<0.001, main effect of LPCAT3 knockdown) but not flotillin1 (P=0.003 main effect of LCPAT3 knockdown, P=0.01 main effect of MβCD) (n=9). (H) Cells were incubated in the presence (0.6 nM) or absence of insulin and were blotted for total or Ser473 phosphorylation of Akt (n=3 Basal, n=6 Insulin). (I&J) C2C12 myoblasts were differentiated into myotubes with either CI-976 or vehicle. (I) Quantification of MC540 fluorescence (*n*=6). (J) Western blot of Thr308 phosphorylation and total Akt in the presence (12 nM) and absence of insulin (*n*=14, P=0.024 main effect of insulin) (A-C&I) Two-tailed t-tests or (E-H&J) 2-way ANOVA with Sidak’s multiple comparisons test were performed. All data are represented as mean ± SEM.

Cholesterol and sphingomyelin are two classes of lipids that are more highly abundant in the detergent-resistant membrane (DRM; i.e. ordered membrane) fraction compared to the detergent-soluble membrane (DSM) fraction (47). Experiments in wild-type C2C12 myotubes indicated that fractions 4-5 have substantial amounts of total lipid (Figure S5A). These fractions were enriched in sphingomyelin and cholesterol which are known to be conducive for more highly ordered membrane (Figure S5B&C), with relatively low abundance of lipids involved in the LPCAT3-mediated reaction (Figure S5D-G). LPCAT3 knockdown did not appear to alter the overall content of lipid in the DRM fraction, nor did it affect enrichment of sphingomyelin and cholesterol (Figure 5B&C). Even though the DRM fraction is known to contain very little protein (48), we detected substantial cav3 in both SC and KD myotubes (Figure 5D). While LPCAT3 deletion did not affect the proportion of cav3 in the DRM fraction, it is noteworthy that elevated cav3 content with LPCAT3 knockdown (Figure 5A) is reflected in the DRM fraction. This is particularly interesting considering there was no enrichment of sphingomyelin or cholesterol in the DRM fraction, which may have been expected given that these lipids induce sequestration of cav3 into caveolae. Consistent with this notion, LPCAT3 deletion was sufficient to elevate lyso-PC in the DRM as well as DRM fractions of the membrane (Figure 5E), with minimal effect on PC species (Figure S5A). Similar results were exhibited with lyso-PE and PE (Figure S5B&C). The saturation index of phospholipids was slightly increased in both DRM and DSM fractions, which may also contribute to the increase in the plasma membrane lipid clustering (Figure S5D). Thus, LPCAT3 deletion promotes the accumulation of lyso-PC in the DRM fraction which may contribute to membrane organization. To test our hypothesis that an increase in membrane organization mediates the insulin-sensitizing effect of LPCAT3 deletion, we incubated C2C12 myotubes with methyl-beta-cyclodextrin (MβCD), a cholesterol-depleting compound that disrupts plasma membrane microdomains (49). Indeed, incubation of cells with MβCD decreased cav3 protein content (Figure 5F) without decreasing the abundance of flotillin-1 (Figure 5G), a protein associated with non-caveolar microdomains. MβCD treatment normalized insulin-stimulated Akt phosphorylation with LPCAT3 deletion to control levels (Figure 5H). These findings are consistent with the notion that LPCAT3 deletion enhances IR signaling by its effect on plasma membrane organization.

CI-976 is a pan lysophospholipid acyltransferase inhibitor (32, 33) that has the ability to disrupt Lands cycle, similar to LPCAT3 deletion. To examine a possibility that the insulin-sensitizing effect of LPCAT3 knockdown is attributable to an unknown function of LPCAT3 outside of the Lands cycle, we studied C2C12 myotubes with or without CI-976. Consistent with our findings with LPCAT3 knockdown (Figure 2E&F), pre-incubation of wild-type C2C12 myotubes with CI-976 robustly decreased phospholipid headgroup packing (Figure 5I). Strikingly, CI-976 also promoted an increase in insulin-stimulated Akt phosphorylation compared to vehicle control (Figure 5J). These evidence support our findings that inhibition of Lands cycle alter plasma membrane property to increase skeletal muscle insulin sensitivity.

## Discussion

Obesity promotes aberrant lipid metabolism in various tissues including skeletal muscle where it dampens its ability to respond to circulating insulin and increase glucose uptake. Studies in model organisms have led to the identification of lipotoxic lipids that might promote insulin resistance in various tissues (50, 51), but some studies were unable to validate these mechanisms in human muscles (52, 53). To gain a global understanding of changes that occur in muscle lipid metabolism with human obesity, we conducted lipidomic analyses on muscle samples from LN and OB subjects. Obesity was associated with decreases in various species of lysophospholipids, an observation that had never been previously reported. Many of these lipids are generated by the enzymes of the Lands cycle, which removes fatty-acyl chains at the sn-2 position of phospholipids to generate lysophospholipids (Lands cycle). We propose a novel mechanism by which obesity accelerates the skeletal muscle Lands cycle to promote insulin resistance.

The acceleration of muscle phospholipid transacylation was apparently driven by increased LPCAT3 expression, likely attributable to diet-induced activation of LXRs and PPARs (22, 54, 55). The inhibition of LPCAT3 enhances insulin signaling at the level of IR to improve skeletal muscle insulin sensitivity. We believe that the insulin-sensitizing effect of Lands cycle inhibition is mediated by its effect on the plasma membrane lipid organization (Figure 6). Consistent with this notion, LPCAT3 deletion and/or CI-976 treatment was sufficient to alter membrane phospholipid packing, GM1-microdomain clustering, cav3 content and lipid composition of detergent-resistant and –soluble membranes. Furthermore, disruption of cholesterol-rich microdomains was sufficient to eliminate the insulin-sensitizing effect of LPCAT3 inhibition. Interventions that interfere with the plasma membrane organization would be predicted to have effects on other cellular events, but the deletion of LPCAT3 did not appear to have an overly adverse effect on skeletal muscle, including mass, fiber-type or force-generating capacity. It would be of substantial interest to pursue implications of altered Lands cycle and/or plasma membrane organization in the context of other cellular events including signaling through other receptor tyrosine kinases.

**Figure 6:**
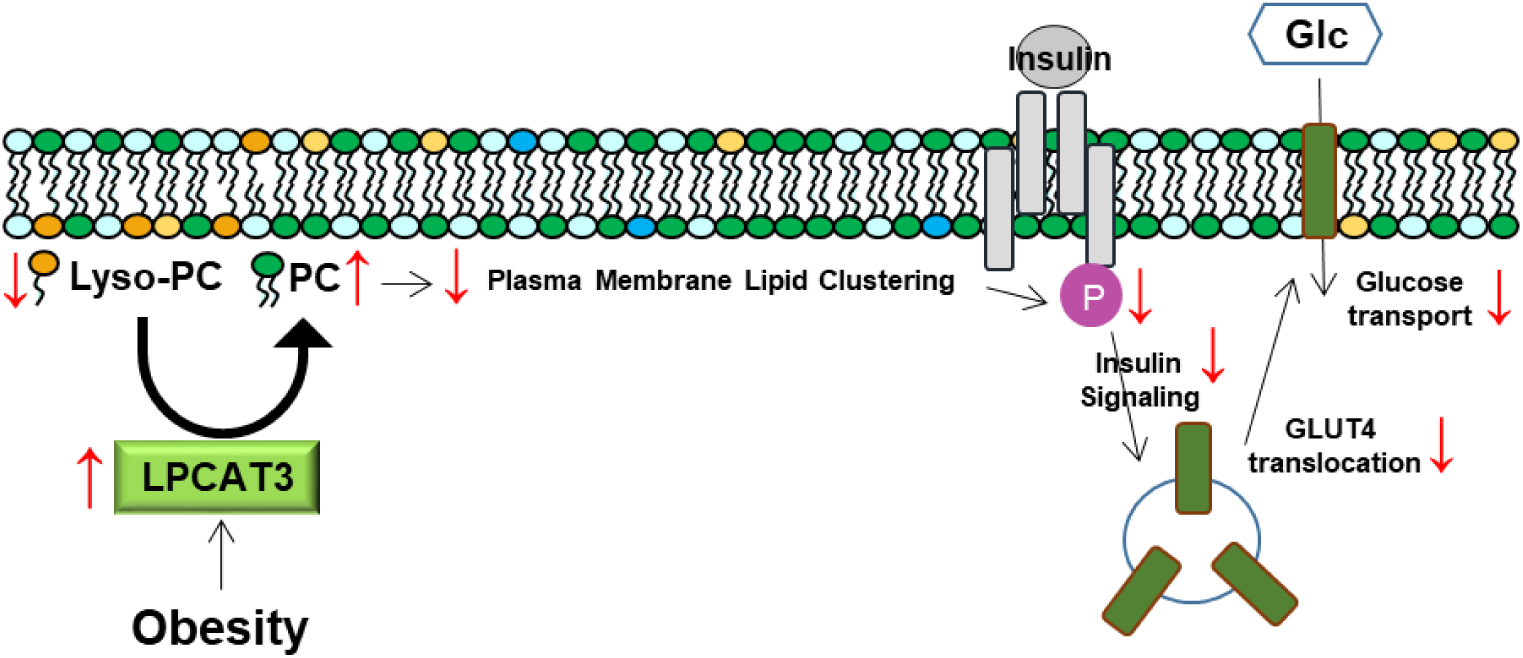
A proposed mechanism of action by which LPCAT3 promotes diet-induced skeletal muscle insulin resistance.

Observations in this study open up a potential opportunity to pharmacologically target this pathway (such as with CI-976) to enhance skeletal muscle insulin sensitivity and improve whole-body glucose homeostasis. It is noteworthy that the current study partly drew its conclusions from lipidomic analyses and loss-of-function studies performed in human samples, suggesting that this mechanism may be directly involved in the pathogenesis of skeletal muscle insulin resistance in human obesity. We are also interested in examining whether obesity induces similar changes in plasma membrane organization of other tissues to promote pathology.

## Methods

### Human Subjects

All participants were prescreened to be free of any known metabolic diseases or heart conditions, nontobacco users, not taking any medications known to alter metabolism, and sedentary. Six lean subjects without diabetes (LN: BMI < 25 kg/m^2^) and six subjects with severe obesity (OB: BMI > 40 kg/m^2^) were studied (all Caucasian females). The subjects were instructed not to exercise for approximately 48h before the muscle biopsy. A fasting blood sample (glucose and insulin) and muscle biopsy from the vastus lateralis were collected. A portion of the biopsy sample was frozen immediately, and another portion was used to isolate primary muscle cells.

### Cell Culture

Primary human skeletal muscle cells (HSkMC) were isolated from fresh muscle biopsies as previously described (15, 56). HSkMC were cultured in growth media containing low glucose DMEM, 10x FBS, 0.5 mg/mL BSA, 0.5 mg/mL fetuin, 10 ng/mL human EGF, 1 µM dexamethasone, and 0.1% penicillin-streptomycin. HSkMC were differentiated in low glucose DMEM, 2% horse serum, 0.5 mg/mL BSA, 0.5 mg/mL fetuin, and 0.1% penicillin-streptomycin. C2C12 myoblasts were grown in high glucose DMEM (4.5 g/L glucose, [+]L-Glutamine; Gibco 11965-092) supplemented with 10% FBS (Heat Inactivated, Certified, US Origin; Gibco 10082-147), and 0.1% penicillin-streptomycin (10,000 U/mL; Gibco 15140122). C2C12 cells were differentiated into myotubes with low glucose DMEM (1 g/L glucose, [+]L-Glutamine, [+]110 mg/L sodium pyruvate; Gibco 11885-084) supplemented with 2% horse serum (Defined; VWR 16777), and 0.1% penicillin-streptomycin. For experiments with CI-976 C2C12 myoblasts were differentiated with either 10 µM of CI-976 or equal volume DMSO (vehicle). For experiments with methyl-beta-cyclodextrin cells were incubated with 10 mM (1320 g/mole) for 1 hour directly dissolved into media. Prior to all experiments cells were serum-starved for 3 hours in low glucose DMEM containing 1% BSA and 0.1% penicillin-streptomycin.

### Quantitative-RT-PCR

Samples were homogenized in TRIzol reagent (Life Technologies, Grand Island, NY) to extract total RNA. 1 μg RNA was reverse transcribed using IScript^TM^ cDNA synthesis kit (Biorad, Hercules, CA). RT-PCR was performed with the Viia^TM^ 7 Real-Time PCR System (Life Technologies, Grand Island, NY) using SYBR® Green reagent (Life Technologies, Grand Island, NY). All data were normalized to ribosomal L32 gene expression and primer sequences are provided (Extended Data Table 2).

### Mass Spectrometry

Global lipidomic analyses for LN and OB HSkMC were performed at the Mass Spectrometry Resource at the Washington University School of Medicine (15). Extracted lipids with internal standards were analyzed with a Thermo Vantage triple-quadrupole mass spectrometer or a Thermo Trace GC Ultra mass spectrometer. Targeted lipidomic analyses for C2C12 myotubes and mouse skeletal muscles were conducted in the Metabolomics Core at the University of Utah (57–59). Extracted lipids with internal standards were analyzed with an Agilent triple-quadrupole mass spectrometer. The quantity of each lipid species was normalized to total lipid content for DRM/DSM experiments or to the total protein content for all others. The phospholipid saturation index was quantified by multiplying the relative abundance of each phospholipid species by the total number of double bonds in the acyl chains of that species.

### Merocyanine 540

Merocyanine 540 (MC540) measurements were taken as previously described (60). In short, skeletal muscle cells (C2C12 and HSkMC) were fully differentiated and 2 million were used for measurements. Cells were washed with Hanks Balanced Salt Solution (HBSS; Gibco 14025092) prior to re-suspension in a cuvette with HBSS. MC540 in DMSO was added at a final concentration of 0.2 µM and after a 10-minute dark incubation, an emission scan was performed ranging from 550-750 nm with fluorescence excitation set at 540 nm on a PTI QuantaMaster 6000 Fluorimeter.

### Lentivirus-Mediated Knockdown of LPCAT3

LPCAT3 expression was decreased using pLKO.1 lentiviral-RNAi system. Plasmids encoding shRNA for mouse LPCAT3 (shLPCAT3: TRCN0000121437) were obtained from Sigma (St. Louis, MO). Packaging vector psPAX2 (ID #12260), envelope vector pMD2.G (ID #12259) and scrambled shRNA plasmid (sc: ID1864) were obtained from Addgene (Cambridge, MA). HEK293T cells in 10 cm dishes were transfected using 50µL 0.1% Polyethylenimine, 200µL 0.15 M Sodium Chloride, and 500 µL Opti-MEM ([+] Hepes, [+] 2.4 g/L Sodium Bicarbonate, [+] L-Glutamine; Gibco 31985) with 2.66 μg of psPAX2, 0.75 μg of pMD2.G, and 3 μg of either scrambled or LPCAT3 shRNA plasmid. After 48 hours, growth media was collected, filtered using 0.22 μm vacuum filters, and used to treat undifferentiated HSkMC or C2C12 cells for 48 hours. To ensure only cells infected with shRNA vectors were viable, cells were selected with puromycin throughout differentiation.

### Western Blot

Whole muscle or cells were homogenized and Western blots were performed as previously described (56). Protein homogenates were analyzed for abundance of phosphorylated(Tyr972)-insulin receptor (Invitrogen: 44-800G), insulin receptor-β (Cell Signaling: 3020S), phosphorylated(Thr308)-Akt (Cell Signaling: 9275S), phosphorylated(Ser472)-Akt (Cell Signaling: 9271L), Akt (Cell Signaling: 9272S), phosphorylated(Thr642)-AS160 (Cell Signaling: 8881), AS160 (Millipore Sigma: 07-741), MyoD (DSHB: D7F2), mitochondrial complexes I-V (Abcam: ab110413), MHC type I (DSHB: A4.840), MHC type IIa (DSHB: SC-71), MHC type IIx (DSHB: 6H1), MHC type IIb (DSHB: BF-F3), MHC neo (DSHB: N1.551), MHC emb (DSHB: BF-G6), Caveolin-3 (BD Biosciences: 610-420), Na/K ATPase (Cell Signaling: 3010S), Flotillin-1 (Cell Signaling: 3253), and actin (Millipore Sigma: A2066).

### Glycogen Synthesis

The glycogen synthesis rate was quantified as previously described (61, 62). Briefly, cells were incubated in media containing D-[U-^14^C] glucose with (12 nM) or without insulin for 2h at 37 °C. Cells were then washed with ice-cold PBS and homogenized for 1h with 0.05% SDS. Part of the lysate was used for a protein assay and the other was combined with 2mg carrier glycogen and incubated at 100 °C for 1h. Ice cold ethanol (100%) was added to the boiled samples prior to overnight rocking at 4 °C. Samples were then centrifuged at 11,000 xG for 15 min at 4 °C to pellet glycogen. Pellets were re-suspended in de-ionized H_2_O and glycogen synthesis was calculated with liquid scintillation.

### Generation of LPCAT3 Skeletal Muscle-Specific Knock Out Mice

Conditional LPCAT3 knock out (LPCAT3cKO+/+) mice were previously generated by flanking exon3 of the *Lpcat3* gene with *loxP* sites (25). LPCAT3cKO+/+ mice were then crossed with tamoxifen-inducible, skeletal muscle-specific Cre-recombinase (HSA-MerCreMer+/-)(34) mice to generate LPCAT3cKO+/+;HSA-MerCreMer−/− (Control; Ctrl) and LPCAT3cKO+/+;HSA-MerCreMer+/− (LPCAT3 Muscle-specific Knock-Out; LPCAT3-MKO). Tamoxifen injected (7.5 µg/g body mass, 5 consecutive days) control and LPCAT3-MKO littermates were used for all experiments. Mice were maintained on a 12 h light/dark cycle in a temperature-controlled room. Prior to all terminal experiments and tissue harvesting, mice were given an intraperitoneal injection of 80 mg/kg ketamine and 10 mg/kg xylazine.

### Glucose Tolerance Test

Intra-peritoneal glucose tolerance tests were performed by injecting 1 mg glucose/g body mass. Mice were fasted for 4 hours prior to glucose injection. Blood glucose was measured prior to glucose injection and 15, 30, 60, and 120 minutes post-injection via tail bleed with a handheld glucometer (Bayer Contour 7151H). In a separate set of experiments, mice were injected with 1 mg glucose/g body mass and blood was taken from the facial vein at the 30-minute time point for insulin quantification.

### Serum Insulin and Glucose Quantification

Blood was collected from the facial vein either prior to anesthesia or at the 30-minute time point of the glucose tolerance test. Blood was then placed at room temperature for 20 minutes to allow for clotting before centrifugation at 2,000 xG for 10 minutes at 4°C. The supernatant (serum) was placed in a separate tube and stored at −80 °C until analysis.

Serum glucose was quantified using a colorimetric assay. A glucose standard curve was generated (Millipore Sigma, G6918) and serum samples were mixed with a PGO enzyme (Millipore Sigma, P7119) and colorimetric substrate (Millipore Sigma, F5803) and measured at OD450 on a plate reader. Serum insulin was quantified using an insulin mouse serum kit (CisBio, 62IN3PEF) using Fluorescence Resonance Energy Transfer on a plate reader (ThermoFisher, Varioskan LUX).

### [^3^H]2-Deoxy-D-Glucose Uptake

*Ex vivo* glucose uptake was measured in the soleus muscle as previously described (63, 64). In brief, soleus muscles were dissected and placed in a recovery buffer (KHB with 0.1% BSA, 8 mM glucose, and 2 mM mannitol) at 37 °C for 10 minutes. After incubation in recovery buffer, muscles were moved to pre-incubation buffer (KHB with 0.1% BSA, 2 mM sodium pyruvate, and 6 mM mannitol) ± 200 µU/mL insulin for 15 minutes. After pre-incubation muscles were placed in incubation buffer (KHB with 0.1% BSA, 9 mM [^14^C]mannitol, 1 mM [^3^H]2-deoxyglucose) ± 200 µU/mL insulin for 15 minutes. Contralateral muscles were used for basal or insulin-stimulated measurements. After incubation muscles were blotted dry on ice-cold filter paper, snap-frozen, and stored at −80 °C until analyzed with liquid scintillation counting.

### Muscle Force Generation

Force generating properties of extensor digitorum longus (EDL) muscles were measured as previously described (65). Briefly, EDL muscles were sutured at each tendon and muscles were suspended at optimal length (L_o_) which was determined by pulse stimulation. After L_o_ was identified muscles were stimulated (0.35 s, pulse width 0.2 ms) at frequencies ranging from 10-200 Hz. Muscle length and mass were measured to quantify CSA (66–68) for force normalization.

### Muscle Immunohistochemistry

Frozen, OCT-embedded hind limb muscle samples (tibialis anterior or EDL) were sectioned at 10µm using a cryostat (Microtome Plus^™^). Following 1h blocking with M.O.M mouse IgG blocking (Vector: MKB-2213), myofiber sections were incubated for 1h with concentrated BA.D5, SC.71, and BF.F3 (all 1:100; DSHB) and laminin (1:200; Millipore Sigma: L9393) in 2.5% normal horse serum. To visualize laminin (for fiber border), myosin heavy chain I (MHC I), myosin heavy chain IIa (MHC IIa), and myosin heavy chain IIb (MHC IIb), slides were incubated for 1h with the following secondaries: AMCA (1:250 Vector: CI-1000), Alexa Fluor 647 (1:250; Invitrogen: A21242), Alexa Fluor 488 (1:500; Invitrogen: A21121) and Alexa Fluor 555 (1:500; Invitrogen: A21426), respectively. Negatively stained fibers were considered myosin heavy chain IIx (MHC IIx). After staining, slides were coverslipped with mounting media (Vector: H-1000). Stained slides were imaged with a fully automated wide-field light microscope (Nikon, Nikon Corp.; Tokyo, Japan) with a 10X objective lens. Images were captured using high sensitivity Andor Clara CCD (Belfast, UK).

### GM-1 Labeling and Imaging

GM-1 clusters were labeled using a Vybrant^®^ Alexa Fluor^®^ 488 Lipid Raft Labeling Kit (ThermoFisher Scientific: V34404) as previously described (69). Briefly, 2 million myotubes were incubated 1mL in ice-cold starvation media with 0.8µg/mL fluorescent cholera toxin subunit B conjugate (CT-B) for 10 minutes. CT-B conjugates were then cross-linked with an anti-CT-B antibody (1:200) in ice-cold starvation media for 15 minutes. Cells were fixed for 1 h at 4 °C in ice-cold 4% paraformaldehyde in PBS in dark. Between each step, cells were washed 2x in ice-cold PBS. Cells were imaged on an Olympus FV1000 confocal microscope (2.5x, HV:600, offset: 30). Images were processed using NIH ImageJ. All images were background subtracted with a rolling ball radius of 50 pixels. Images were blindly scored by S.R.S and K.F. as exhibiting clustering of microdomains or non-clustering. Images were then subjected to color thresholding using the Otsu method (70, 71) (designed for thresholding images for cluster analyses) and made binary. A particle analysis of all particles that were >0.1µm^2^ was performed to determine the average cluster size for each cell that was imaged (72). For each experiment, 35-50 cells per group were analyzed and the median was taken as a representative of that experiment.

### Detergent-Resistant Membrane Isolation

Detergent-resistant membranes (DRM) and detergent-soluble membranes (DSM) were isolated as previously described (72). Briefly, 2×15 cm plates of cells were scraped in ice-cold PBS and then pelleted. Cells were re-suspended in 1mL of cold homogenization buffer (Mes-buffered saline [MBS], 1% Triton-X wt/v, and protease and phosphatase inhibitor) and passed through a 23-gauge needle 6 times before incubating at 4 °C for 30 minutes. MBS was added to the homogenate until a volume of 2.5 mL was reached then mixed with 2.5 mL of 90% sucrose in MBS and 4mL of this mixture was added to an ultracentrifuge tube (Beckman Coulter 344061). A sucrose gradient was generated by adding 4 mL 35% sucrose followed by 4mL of 5% sucrose. Samples were then centrifuged at 100,000 xG at 4 °C for 20 hours in a swinging bucket rotor (Beckman L8-M Ultracentrifuge, SW28 Rotor).

### Statistics

Statistical analysis was performed using Prism 7 software (GraphPad). Student’s t-tests were performed with data composed of 2 groups and 2-way ANOVA for multiple groups followed by Sidak’s multiple comparison test. All data are Mean±SEM and statistical significance was set at P<0.05.

### Study Approval

The experimental protocol was approved by the Internal Review Board for Human Research at East Carolina University. Informed consent was obtained prior to inclusion in the study.

Animal experiments were approved by the University of Utah Institutional Animal Care and Use Committee.

## Author contributions

P.J.F. and K.F. contributed to study concept design and wrote the manuscript. J.A.H. performed human muscle biopsies. J.M.J. contributed to study concept and design and data analysis. X.R. and P.T. developed LPCAT3 conditional knock-out mice. J.A.M., J.E.C., H.S., and J.T. performed mass spectrometry analyses. P.J.F., K.F., and S.R.S. performed analyses of the physical properties of phospholipid membranes. A.R.P.V. and P.J.F. performed analysis of muscle force production. P.S. performed muscle histology measurements. P.J.F. performed all biochemical assays and metabolic phenotyping measurements. K.C.K. and A.J.L. performed correlation analyses with 106 mouse strains. J.A.H., P.T., J.T., J.E.C., and S.R.S. edited the manuscript.

## Acknowledgments

This work was supported by NIH grants DK107397, DK109888, AG063077 (to K.F.), HL030568, HL030568, HL136618 (to P.T.), AT006122, HL123647 (to S.R.S.), HL028481, HL030568 (to A.J.L.), and DK056112 (to J.A.H.), American Heart Association 18PRE33960491 (to A.R.P.V.), American Heart Association 19PRE34380991 (to J.M.J.), and the Larry H. and Gail Miller Family Foundation (to P.J.F). The University of Utah Metabolomics, Mass Spectrometry, and Proteomics core is supported by S10 OD016232, S10 OD021505, and U54 DK110858. The Washington University Biomedical MS Resource is supported by United States Public Health Services grants P41-GM103422, P30-DK020579, and P30-DK056341.

---

**Figure S1:**
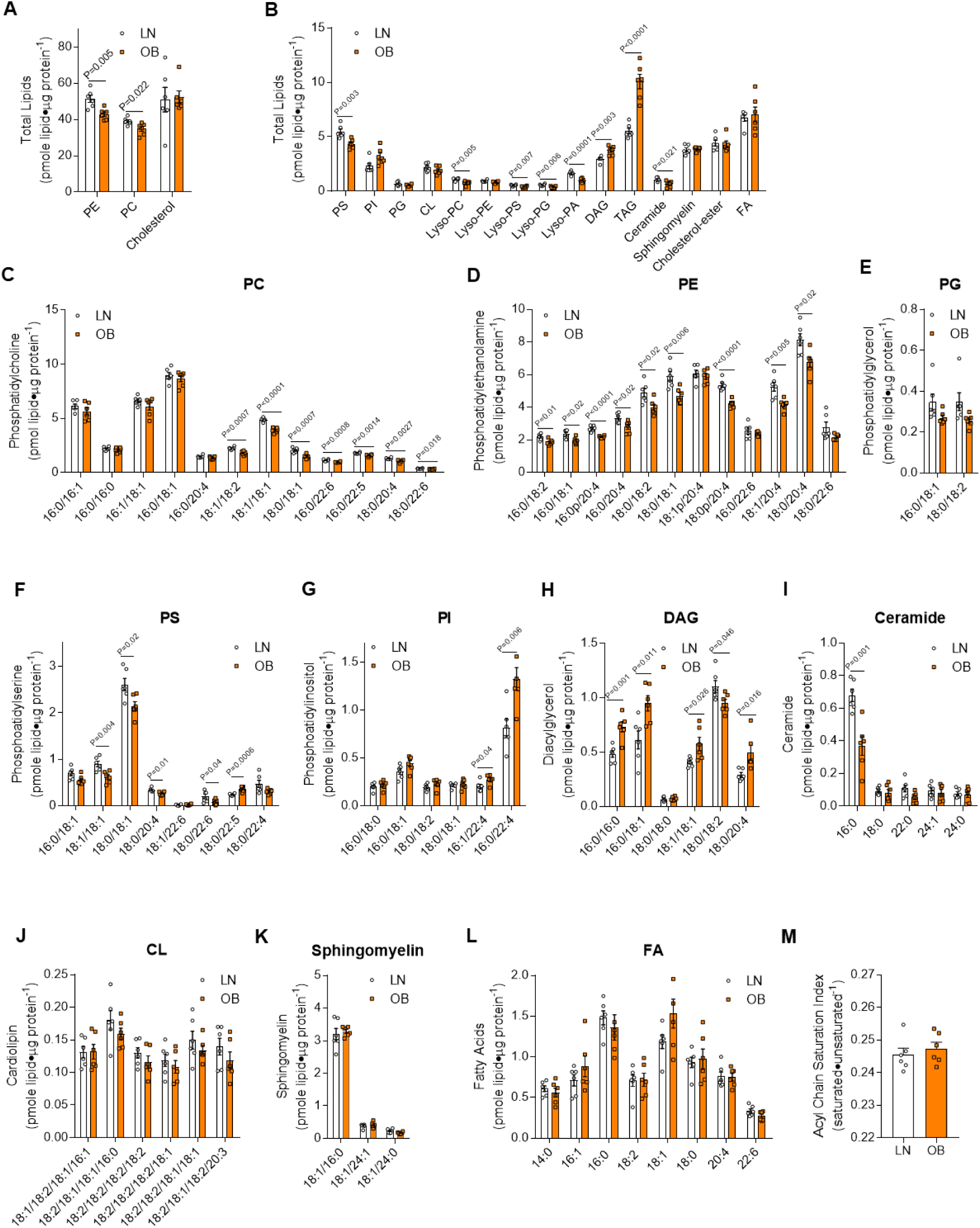
Lipid quantification in LN and OB HSkMC. (A-M) Muscle biopsies were taken from LN or OB human subjects and primary skeletal muscle cells were isolated and differentiated. Quantification of (A&B) total lipids by class, and species of (C) phosphatidylcholine (PC), (D) phosphatidylethanolamine (PE), (E) phosphatidylglycerol (PG), (F) phosphatidylserine (PS), (G) phosphatidylinositol (PI), (H) diacylglycerol (DAG), (I) ceramide, (J) cardiolipin (CL), (K) sphingomyelin, and (L) fatty acid (FA). (M) Quantification of the acyl chain saturation index of all detectable phospholipids. (*n*=6). Two-tailed t-tests were performed for all analyses. All data are represented as mean ± SEM.

**Figure S2:**
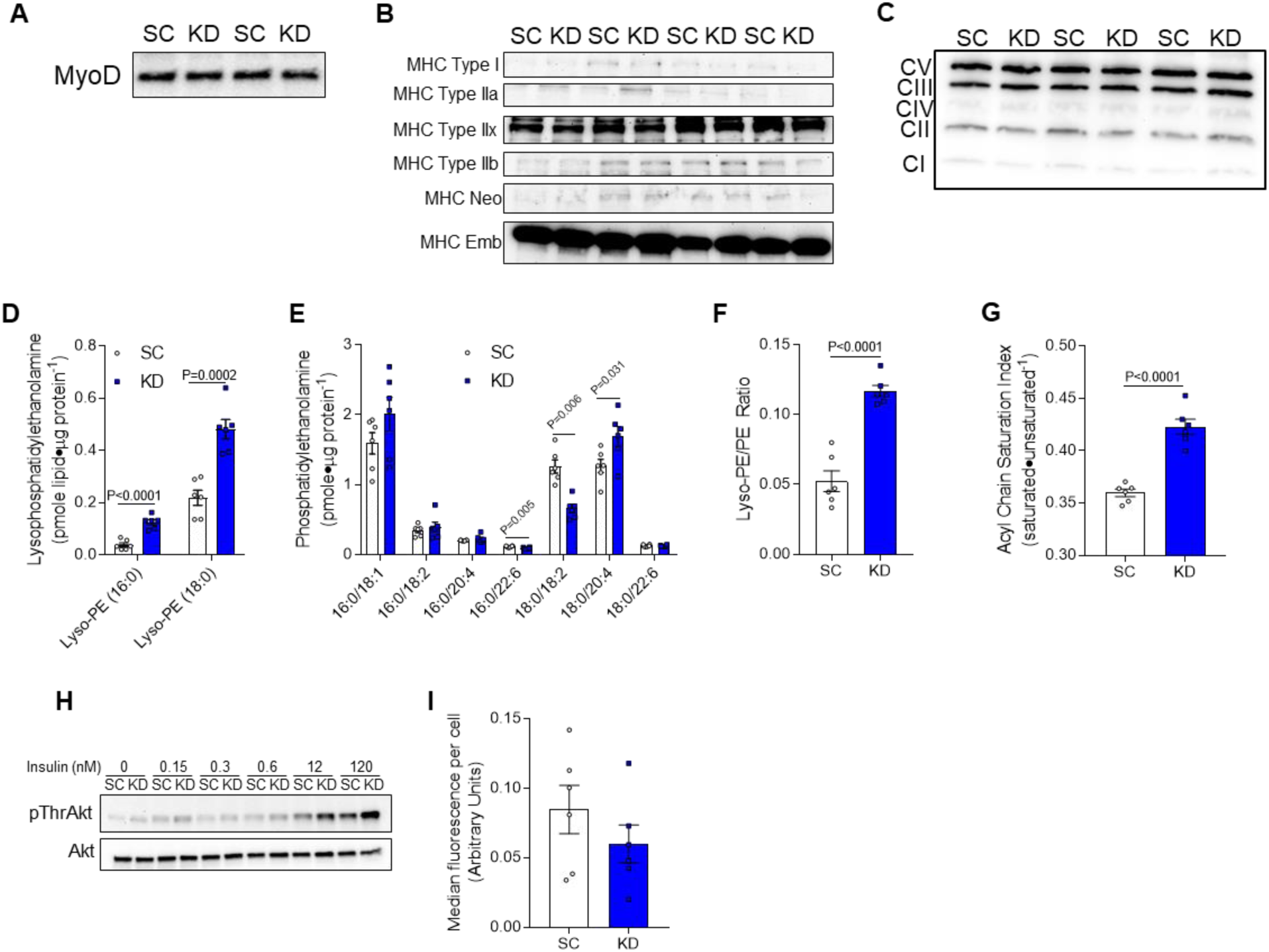
Knockdown of LPCAT3 in C2C12 myotubes. C2C12 myoblasts were infected with shRNA generating lentiviruses targeting scrambled (shScrambled; SC) or LPCAT3 (shLPCAT3; KD) to decrease LPCAT3 expression and cells were differentiated into myotubes. (A-C) Western blot in SC and KD cells probing for (A) MyoD, (B) myosin heavy chain isoforms, and (C) complexes I-V of the electron transport chain. (D-G) Lipids were extracted in SC and KD myotubes for quantification of (D) lyso-PE, (E) PE species, (F) total lyso-PE/PE, and (G) acyl chain saturation index of phospholipids (*n*=6). (H) Thr308 phosphorylation and total Akt from cells incubated (10 min) with various concentrations of insulin. (I) GM-1 microdomains were labeled with GFP and cross-linked to induce patching in SC and KD C2C12 myotubes. Total fluorescence in each cell was measured across 6 separate experiments (*n*=35-50/experiment) and the median of each experiment was used as a representative. Two-tailed t-tests were performed. All data are represented as mean ± SEM.

**Figure S3:**
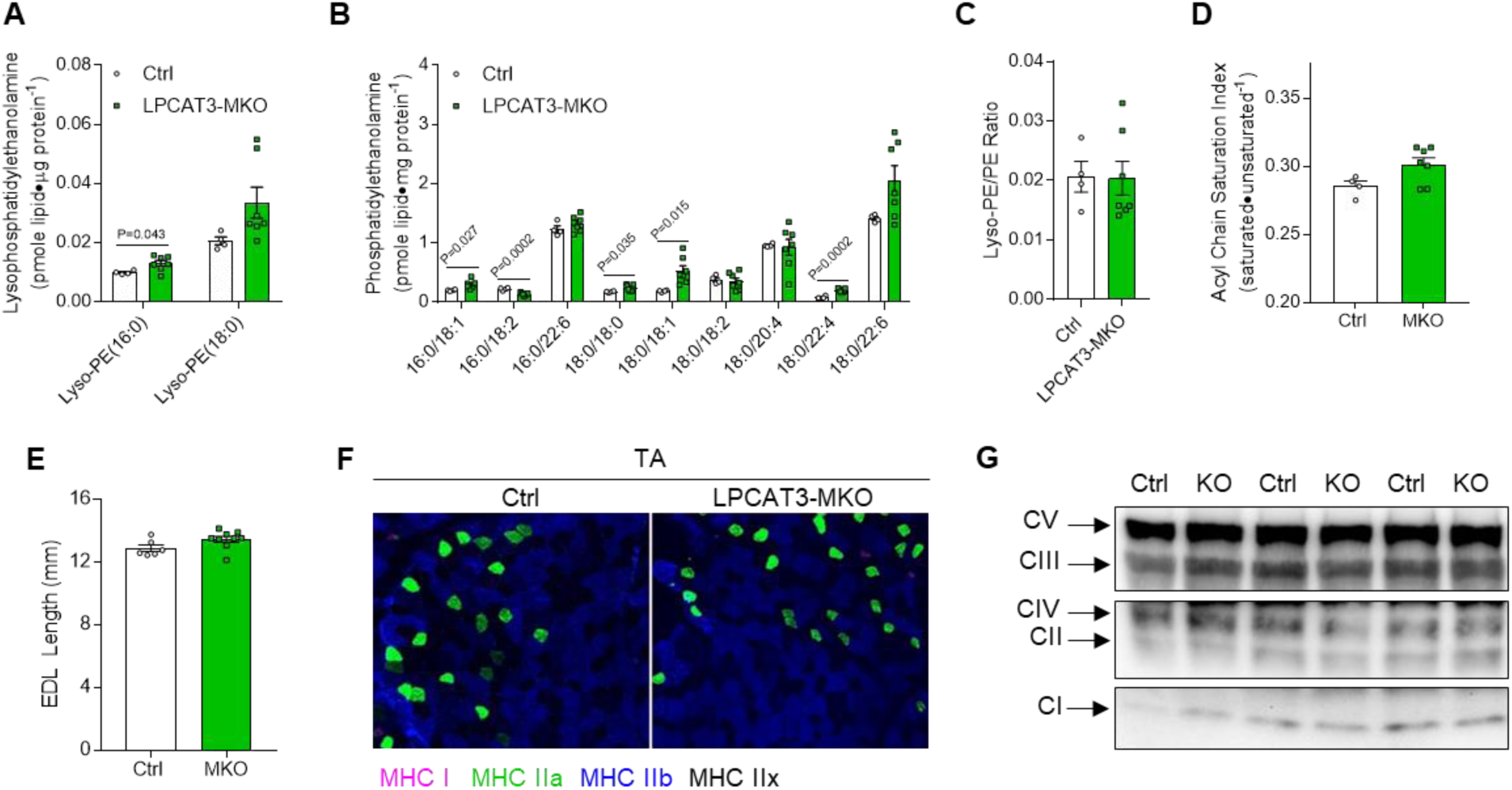
Additional data on muscles from HFD-fed Ctrl and LPCAT3-MKO mice. (A-D) Lipids were extracted from gastrocnemius muscles of Ctrl and LPCAT3-MKO mice for analysis. Quantification of (A) lyso-PE species, (B) PE species, (C) total lyso-PE/PE, and (D) phospholipid acyl chain saturation index (Ctrl *n*=4, MKO *n*=7). (E) Muscle lengths of extensor digitorum longus (EDL) muscles (Ctrl *n*=6, MKO *n*=9). (F) Skeletal muscle fiber-type (MHC I: pink, MHC IIa: green, MHC IIb:blue, and MHC IIx: negative) of tibialis anterior (TA) muscles. (G) Measurement of complexes I-V of the electron transport chain in TA muscles from Ctrl and LPCAT3-MKO mice. (A-E) Two-tailed t-tests. All data are represented as mean ± SEM.

**Figure S4:**
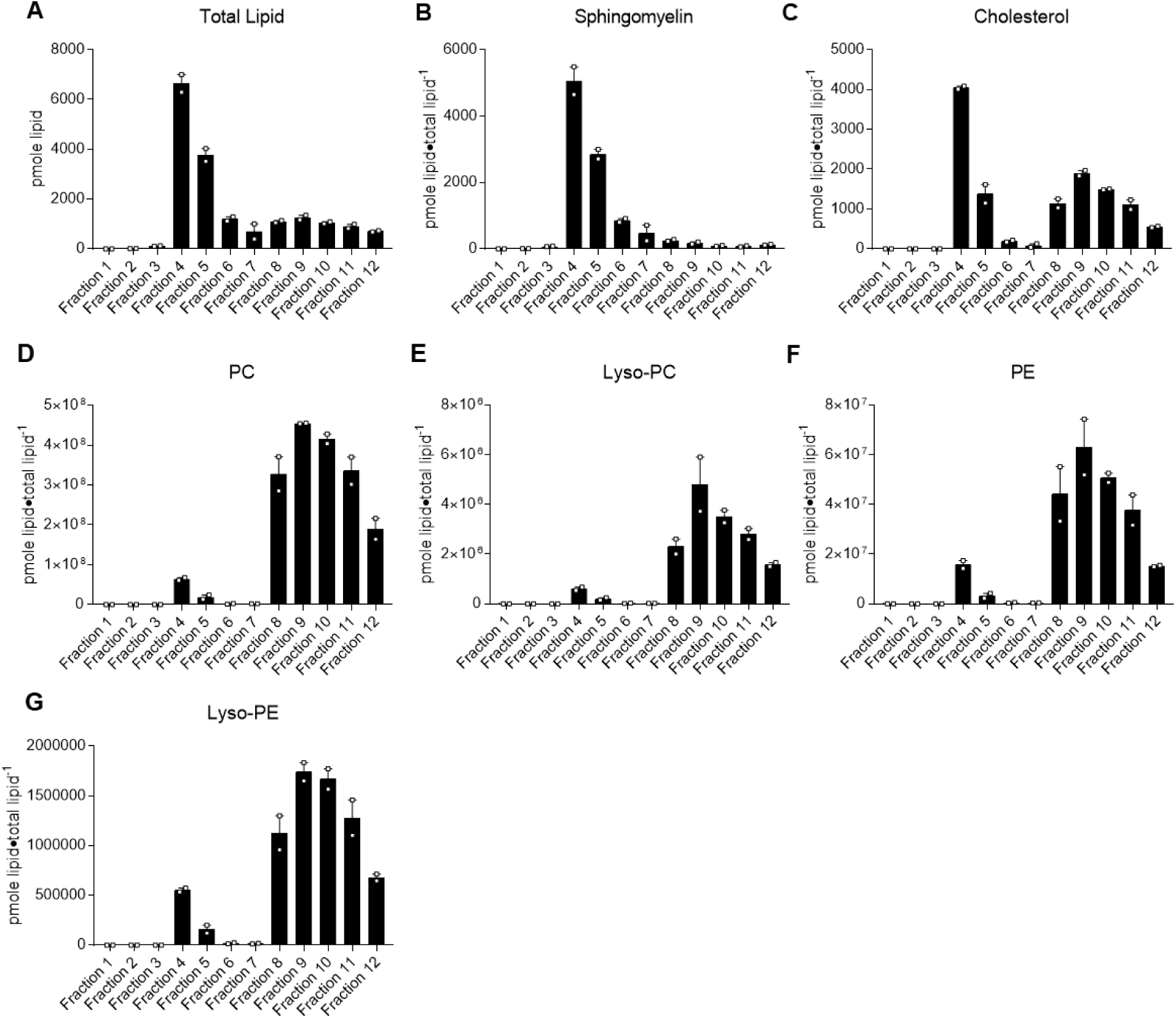
Lipid contents of detergent-resistant and detergent-soluble membrane fractions. (A-G) Wild type C2C12 myotubes were suspended in a sucrose gradient and purified via ultracentrifugation to separate detergent-resistant membrane (DRM) fractions from detergent soluble membrane (DSM) fractions. After ultracentrifugation fractions were analyzed for (A) total lipid content, (B) sphingomyelin, (C) cholesterol, (D) PC, (E) lyso-PC, (F) PE, and (G) lyso-PE (*n*=2). All data are represented as mean ± SEM.

**Figure S5:**
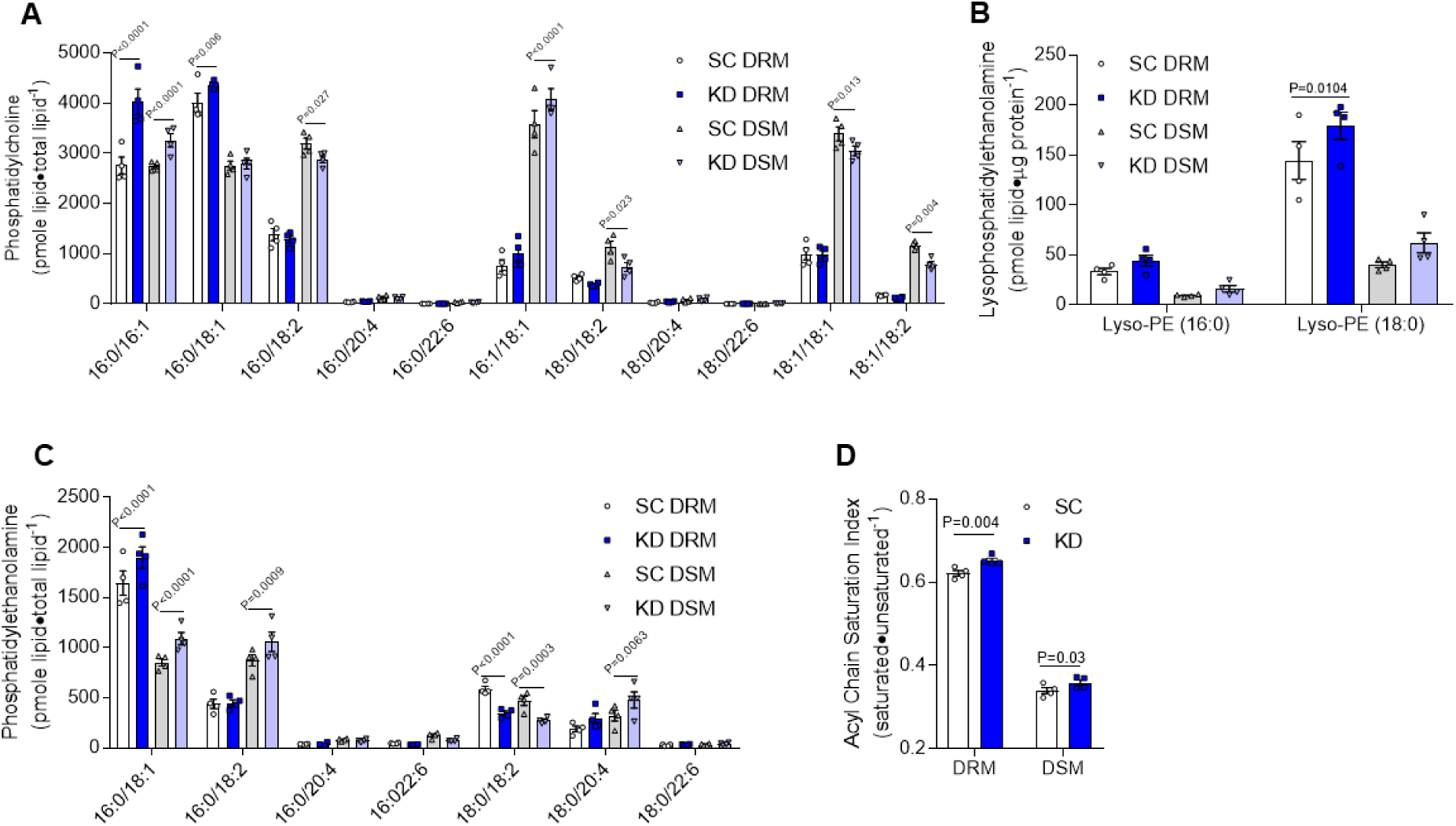
Plasma membrane microdomains and phospholipid composition in membrane fractions with LPCAT3 inhibition. (A-D) SC and KD C2C12 myotubes were suspended in a sucrose density gradient and purified with ultracentrifugation. (A) PC, (B) lyso-PE, (C) PE species, and (D) acyl chain saturation index of all phospholipids were quantified in DRM and DSM fractions (*n*=4). (D) Two-tailed t-tests or (A-C) 2-way ANOVA with Sidak’s multiple comparisons test were performed. All data are represented as mean ± SEM.

**Table S1:** Subject characteristics of lean insulin-sensitive (LN) and obese insulin-resistant (OB) subjects (*n*=6/group).

|  | LN | OB | p |
| --- | --- | --- | --- |
| Age (year) | 30.2±3.5 | 35.8±3.1 | 0.25 |
| Height (cm) | 162.4±2.8 | 166.9±2.6 | 0.26 |
| Weight (kg) | 62.9±2.4 | 126.1±8.0 | <0.0001 |
| BMI ($\text{kg}\cdot\text{m}^{-2}$) | 23.85±0.67 | 45.0±1.85 | <0.0001 |
| Glucose ($\text{mg}\cdot\text{dL}^{-1}$) | 81.3±1.0 | 92.3±4.3 | 0.031 |
| Insulin ($\mu\text{U}\cdot\text{mL}^{-1}$) | 6.4±1.3 | 15.9±1.3 | 0.0004 |
| HOMA-IR | 1.29±0.27 | 3.6±0.44 | 0.001 |
| Cholesterol ($\text{mg}\cdot\text{dL}^{-1}$) | 177.7±13.0 | 177±12.9 | 0.97 |
| HDL ($\text{mg}\cdot\text{dL}^{-1}$) | 55.2±2.3 | 47.0±2.9 | 0.050 |
| LDL ($\text{mg}\cdot\text{dL}^{-1}$) | 105.2±11.9 | 109.5±8.6 | 0.78 |
| Triglycerides ($\text{mg}\cdot\text{dL}^{-1}$) | 86.8±19.8 | 102.7±23.7 | 0.62 |

**Table S2:**
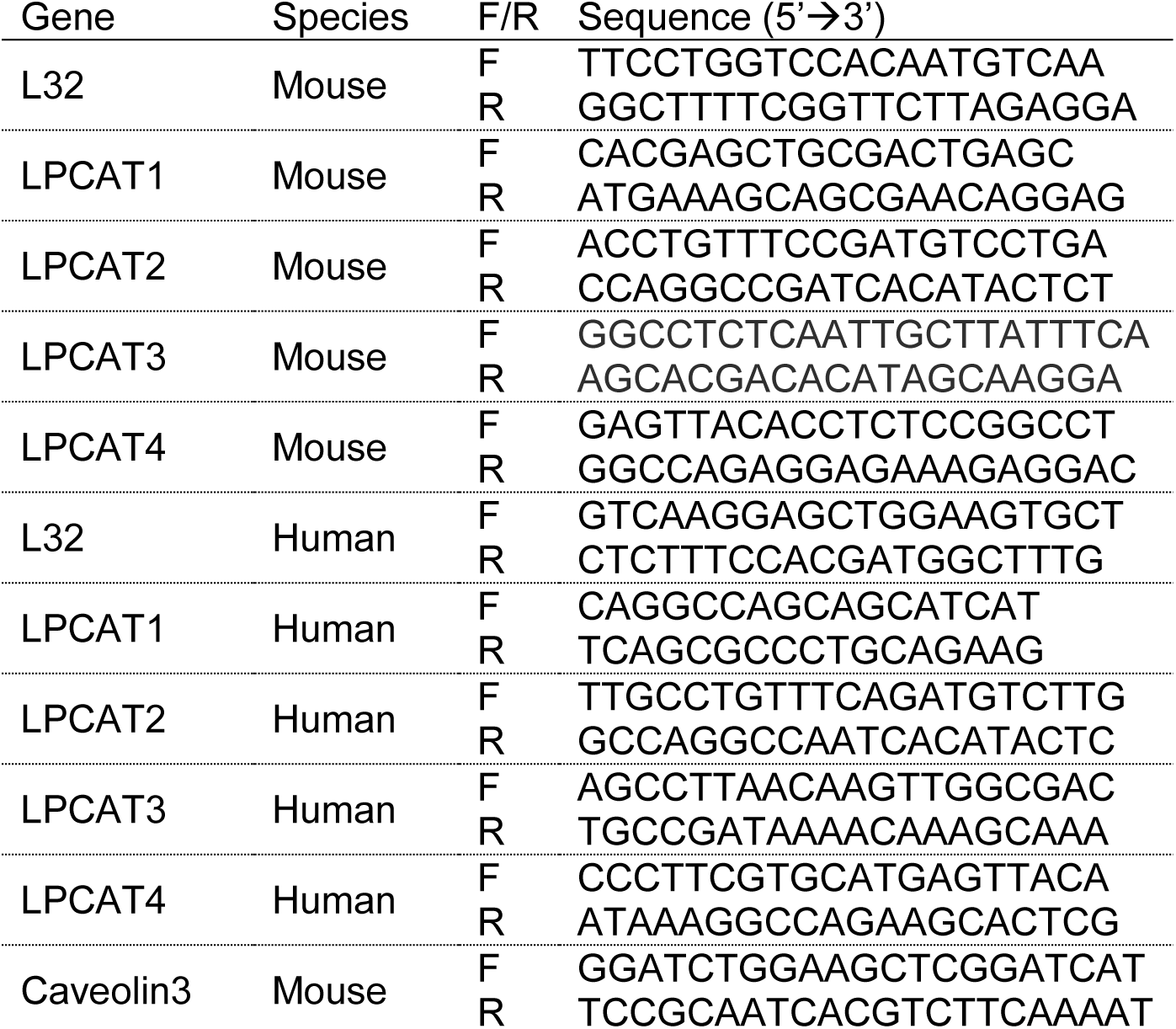
Primers used for quantitative-RT-PCR.

